# Climate change has affected the spillover risk of bat-borne pathogens

**DOI:** 10.1101/2022.09.06.506814

**Authors:** P Van de Vuurst, H Qiao, D Soler-Tovar, LE Escobar

## Abstract

Bat-borne viruses are a threat to global health and have in recent history had major impacts to human morbidity and mortality. Examples include diseases such as rabies, Ebola, SARS-Cov-1, and SARS-Cov-2 (COVID-19). Climate change could exacerbate the emergence of bat-borne pathogens by affecting the distribution and abundance of bats in tropical ecosystems. Here we report an assessment of historical climate and vampire bat occurrence data for the last century, which revealed a relationship between climatic variation and risk of disease spillover triggered by changes in bat distributions. This report represents one of the first examples of empirical evidence of global change effects on continental patterns of bat-borne pathogen transmission risk. We therefore recommend that more research is necessary on the impacts of climate change on bat-borne pathogen spillover risk, and that climate change impacts on bat-borne disease should be considered in global security initiatives.

**Highlights:**

- Bat-borne viruses are a threat to global health and include diseases such as rabies, Ebola, SARS-Cov-1, and SARS-Cov-2 (COVID-19).
- Climate change could exacerbate the emergence of bat-borne pathogens by affecting the distribution and abundance of bats.
- Here we report an assessment of historical climate and vampire-bat occurrence data for the last century, which reveals a relationship between climatic variation and risk of disease spillover triggered by changes in bat distributions.

## 1.0 Introduction

Human manipulation of landscapes can modify the natural interaction between wildlife and their pathogens, which can result in the emergence or re-emergence of infectious diseases (David M. Morens et al., 2004). Interactions among populations of humans, livestock, and wildlife associated with these changes can facilitate pathogen spillover (i.e., the transmission of pathogens from one species to another) (Plowright et al., 2017) (Figure 1). A series of factors have been proposed as facilitators of pathogen spillover from wildlife to humans, including urbanization (Gibb et al., 2020), agricultural practices (Zhou et al., 2018), bushmeat consumption (Pernet et al., 2014), and wet market presence (Webster, 2004). Nevertheless, the interconnected and intersectional nature of the factors that facility or prevent spillover can make this phenomenon difficult to study (Plowright et al., 2017). The public health burden of spillover is extensive, with many zoonotic pathogens having heavy impacts on human morbidity and mortality (Christou, 2011; Smith et al., 2009; Zhou et al., 2018). Nevertheless, one area of scientific uncertainty and concern is the potential impact of climate change on zoonotic disease emergence (Carlson et al., 2022).

**Figure 1:**
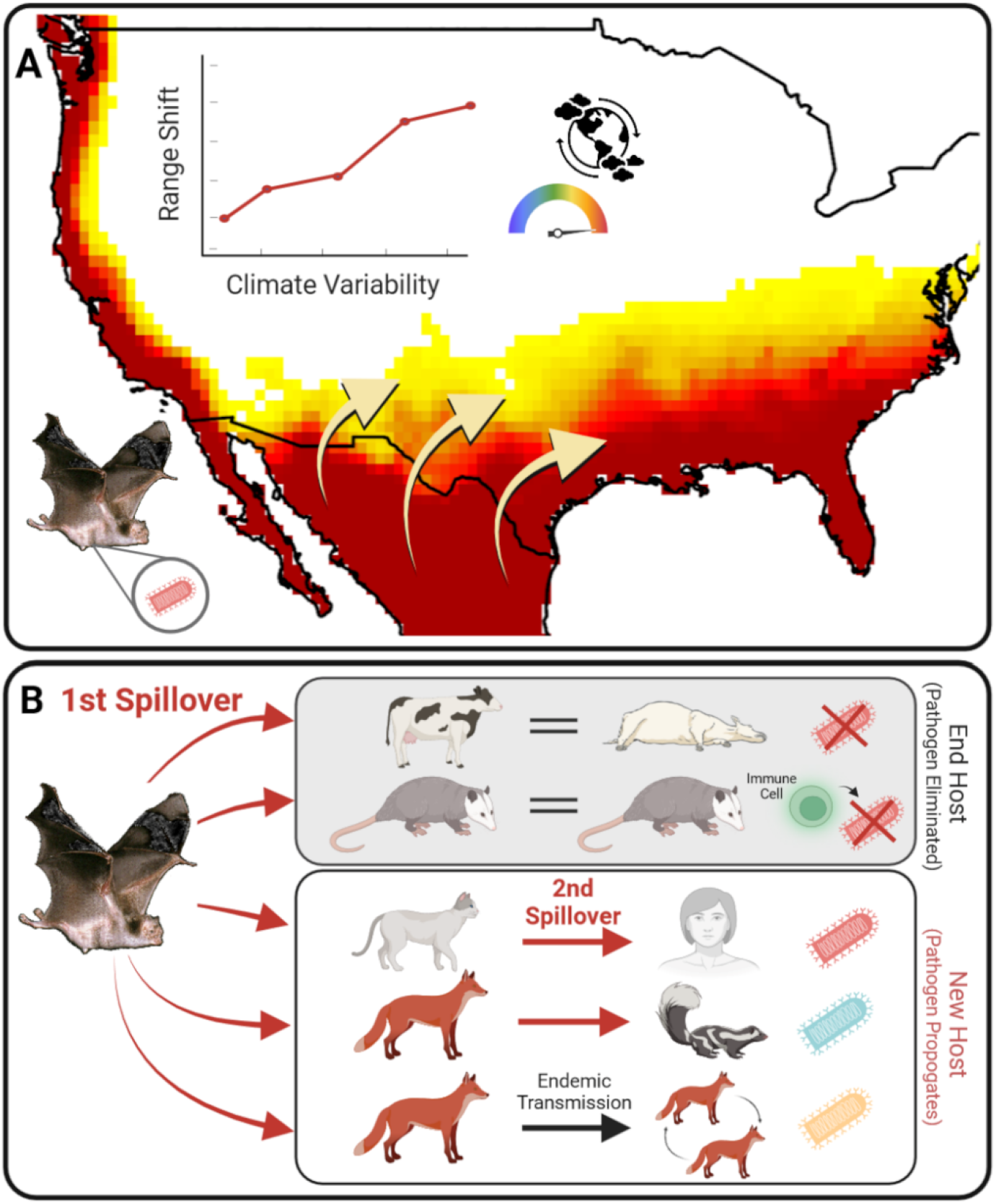
Possible bat-borne pathogen spillover transmission triggered by climate change driven distributional change. **(A)** The common vampire bat (*Desmodus rotundus*) has steadily shifted its range northward in responses do climate change based (Supplementary Methods 2) and could move into more temperate regions of the United State. As a result, **(B)** rabies virus could be transmitted from an infected bat (i.e., Reservoir) to other animal species (i.e., spillover), which may or may not maintain transmission (i.e., end host in grey box vs. new host in white box), which is the prelude for disease emergence and outbreaks. Spillover can also occur from animals to humans (i.e., zoonosis). Figure made using BioRender.

Climate change has been shown to trigger shifts in the distribution of vector-borne disease (Siraj et al., 2014). More simply put, this means that vectors such as mosquitoes have shifted their geographic distributions in response to changes in temperature and humidity, and as a result have invaded new areas. Thus climate change acts as a change agent, modifying the spatial, seasonal, and interannual occurrence of infectious diseases (e.g., the extension of malaria into regions at higher elevations (Siraj et al., 2014), increased incidence and geographic spread of Valley fever (Gorris et al., 2019), or increased abundance of *Vibrio* bacteria, the causative agent of diseases such as Cholera (Vezzulli et al., 2016). Recent research on climate change impacts on disease has largely focused on vector-borne diseases such as Dengue virus (Messina et al., 2019). These studies, however, have generally been based on future climate change projections, rather than retrospective empirical data, which could hamper progress in global infectious disease preparedness. Furthermore, little research has been conducted on directly transmitted diseases within the context of climate change, presenting a possible blind spot in the current understanding of climate change impacts.

## 2.0 Bats and Emerging Infectious Diseases

For decades the scientific community has recognized the association between bats and emerging diseases (Brook & Dobson, 2015). Many infectious agents such as Hendra virus (Eaton et al., 2006; Mahalingam et al., 2012), Ebola virus(Leroy et al., 2005), Nipah virus (Chua et al., 2000), SARS-Cov-1 (Li et al., 2005), MERS (Memish et al., 2013), and SARS-Cov-2 (Latinne et al., 2020) have originated from viral spillover from bats to intermediate animal hosts (Letko et al., 2020) (Figure 1). As such the continue monitoring of bat-borne pathogens is necessary to anticipate future epidemics, especially within the context of climate change. Any change in the abundance, composition, or distribution of bat communities could present an opportunity for novel infectious disease emergence in new geographic areas or animal species. There are no simple solutions for the problem of bat-borne emerging infectious diseases, as bats serve as important pollinators for many plant species, and culling campaigns have proven to be ineffective in stopping the spread of bat-borne infections (Streicker et al., 2012). Understanding how and to what extent climate change may influence bat ecology is therefore of critical importance for the understanding and prediction of emerging bat-borne diseases.

## 3.0 Bat-borne rabies in the Americas as an empirical case

Rabies is one of the oldest infectious diseases to affect humans in recorded history and one of the most lethal of all zoonotic diseases (Velasco-Villa, Mauldin, et al., 2017). Approximately 50,000 recorded human deaths due to rabies are reported annually despite vaccination efforts, with many of these deaths occurring in at-risk populations (Hampson et al., 2015). Rabies also has precipitous impacts to livestock, domesticated pets, and wildlife as well (Meske et al., 2021). Bats are a key reservoir of rabies virus in the Americas (Meske et al., 2021). The common vampire bat (*Desmodus rotundus*), a sanguivorous species in the Phyllostomidae family, is considered to be the main species responsible for transmitting rabies to other species in Latin America (Meske et al., 2021). As a model disease system, vampire bat transmitted rabies is one of the most well understood and documented directly-transmitted diseases to still impact humans and animals in tropical and sub-tropical regions of the Americas. In this region, rabies is commonly transmitted from *D. rotundus* to cattle. Indeed, the bulk of cattle rabies in this region is caused by vampire bats, with canine rabies having been almost eliminated (Velasco-Villa, Escobar, et al., 2017; Velasco-Villa, Mauldin, et al., 2017). Recent years have even seen an increase in rabies in Latin America, with over 1,500 animal cases being reported in 2021 alone (Pan American Health Organization, 2022). Furthermore, according to data from the Pan American Health Organization, rabies transmission from vampire bats to cattle has increased, ranging from 1,000 to 12,500 percent in some countries from the 1970’s to the 2010’s (Figure 2). The largest increases in cases occurred in Peru, Mexico, Ecuador, and Brazil (Figure 2).

**Figure 2.**
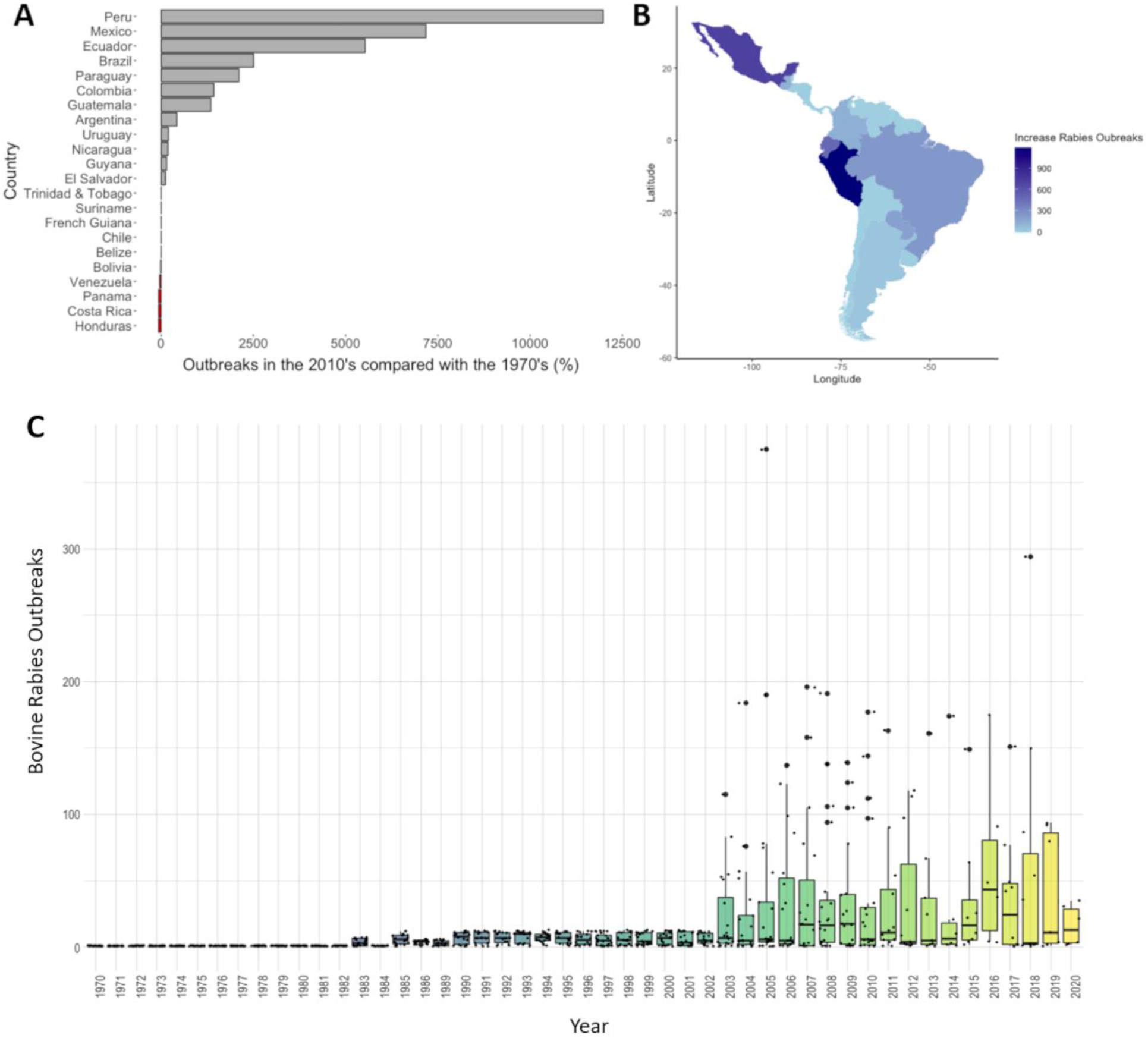
Bat-borne rabies in livestock in Latin America 1970’s vs. 2010’s. **(A)** Percentage change in rabies outbreaks in cattle for Latin American countries. (**B)** Geographic distribution of changes in the number of bat-borne rabies outbreaks in cattle (Supplementary Methods S1). **(C)** Box plots of bovine rabies outbreaks per year (each point represents a country).

### 3.1 Retrospective climate change impacts on vampire bats

Debate still exists regarding the potential response of vampire bats to climate change, including whether or not the species could extend its range into warm-temperate regions in higher latitudes or elevations in the future (Piaggio et al., 2017). Using historic records of *D. rotundus* occurrence from the last century (1900-2020) (Van de Vuurst et al., 2021) and climate data from the Climatic Research Unit gridded Time Series database (CRU TS) version 4.04 database (Harris et al., 2020) we assessed *D. rotundus* distributions in response to climatic variation using machine-learning ecological niche modeling (Supplementary Methods S2). We found no significant change in total area of *D. rotundus* distribution during this time period. Nevertheless, we found that *D. rotundus’* geographic range has moved northward based on this analysis (Figure 3). These results indicates that vampire bats have moved steadily into northern Mexico at an average rate of 7.93 km per year (Supplementary Methods S2), and very well could extend their range into the continental United States if this pattern continues. We also found areas of uncertainty for range expansion, including highlands, such as the Andes Mountains, and in temperate portions of the United States where *D. rotundus* establishment is uncertain but could be vulnerable to a *D. rotundus* associated rabies expansion (Figure 3). The variable that most influenced the range shift in *D. rotundus* distribution was temperature variability (i.e., standard deviation), a variable closely linked with changes in climate.

**Figure 3:**
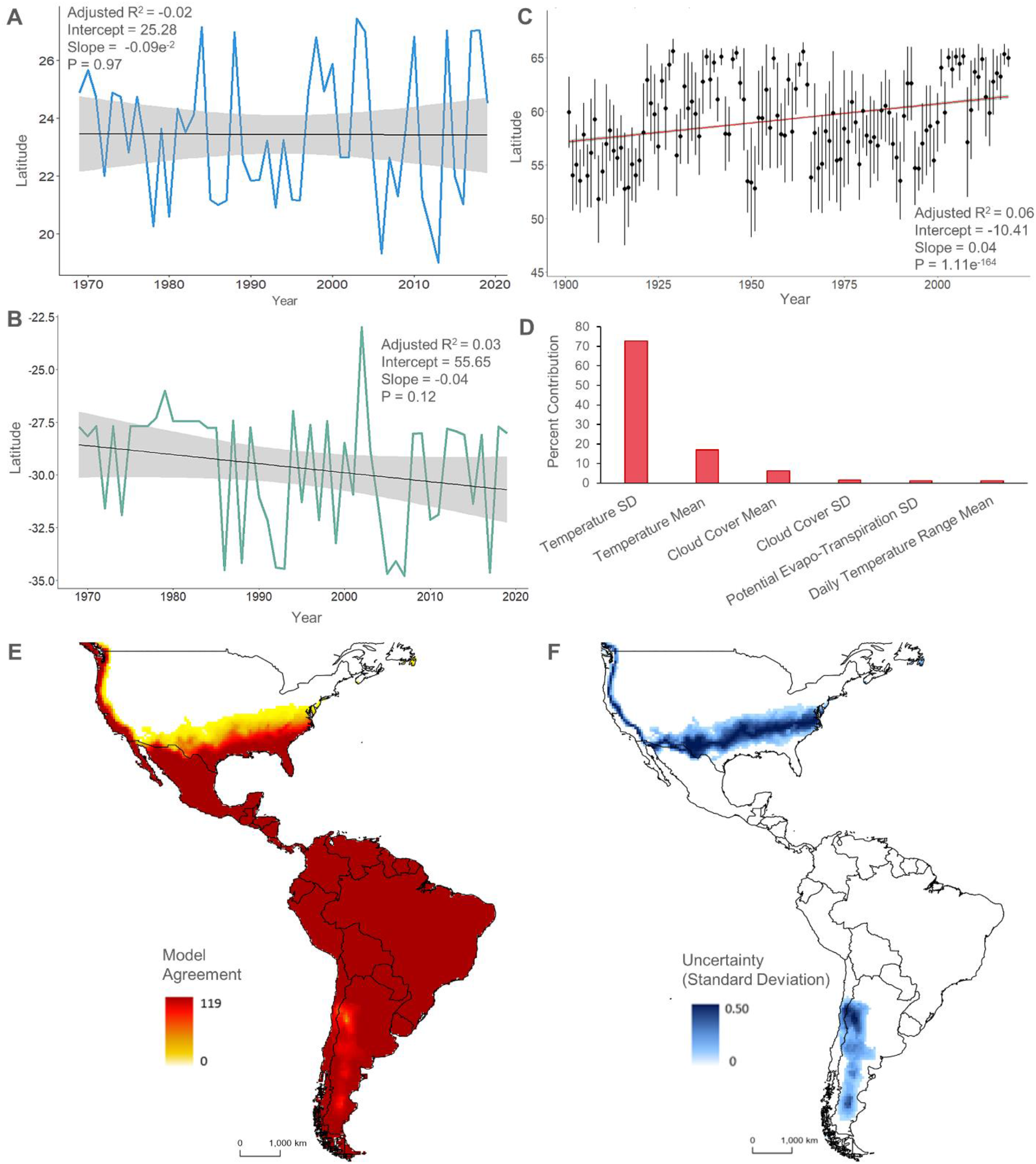
Range shift of *Desmodus rotundus* due to climate change. Evidence of *D. rotundus* range shifts, both North (**A**) and South (**B**) in the last 50 years based on species reported occurrence. (**C**) Predicted northward trend (based on ecological niche models) of D. rotundus range across the last century. (**D**) Estimates of relative percent contributions of the environmental variables on the distribution of *D. rotundus*. Note that temperature variables (standard deviation and mean), and cloud cover contribute most to the model. (**E**) Model ensemble of *D. rotundus* distributions from 1901 to 2019. (Darker colors indicate higher agreement in models regarding *D*.*rotundus* distributions during the 119-year period). (**F**) Uncertainty map revealing areas with higher uncertainty (darker colors of blue) with regard with the potential areas of *D. rotundus* expansion. Note uncertainty in the southern United States and portions of the Andes Mountains.

## 4.0 Conclusion

Results from this research report indicate that bat-borne rabies outbreaks in cattle have risen in the last 40 years and reflect an increase in spillover events in the American continents (Figure 2). Furthermore, a century-long, empirical dataset of *D. rotundus* occurrence has also demonstrated that climate change has impacted the distribution of a virus reservoir across time (Figure 2). Climate change, therefore, has directly impacted the risk of rabies incidence and pathogen spillover by increasing the presence of the virus reservoir in temperate areas of the Americas, putting humans, wildlife, and livestock at risk for infection. As such, this evidence provides an empirical, retrospective example of climate change driven range shifts of a bat reservoir, which until now has been limited to future climate simulations (Carlson et al., 2022). This evidence indicates that infectious disease emergence and viral spillover will increase under future climate scenarios. As the global climate continues to change, the risk of future bat-borne pandemics will remain relevant for global health security as a result. The current uncertainties surrounding future climate change impacts on emerging infectious disease could be mitigated by developing retrospective measurements of the effects of climatic variation on disease transmission. Understanding the underlying causes of disease transmission, and the likely ripple effects of climate change upon bat-borne pathogens should be considered in a global health security agenda to better prepared for the next pandemic.

## Acknowledgments

The authors would like to acknowledge the efforts of Natalie Brown for her assistance with the completion of this work. This study was supported by the National Science Foundation award: Human-Environment and Geographical Sciences Program 2116748.

