## Supplemental Methods for "Climate change has affected the spillover risk of bat-borne pathogens"

### Supplementary Materials

#### 1.0 Supplementary Methods 1

Data on the presence of bovine rabies cases transmitted by *Desmodus rotundus* were collected from the Regional Information System for the Epidemiological Surveillance of Rabies (SIRVERA) (PANAFTOSA, 2021). SIRVERA is a database for the prevention of rabies in the Americas, where the countries of the American continents report the presence of rabies on a monthly basis (Belotto, Leanes, Schneider, Tamayo, & Correa, 2005; Benavides et al., 2019; Del Rio Vilas et al., 2017; Freire de Carvalho et al., 2018; Rysava et al., 2020; Seneschall & Luna-Farro, 2013; Velasco-Villa et al., 2017; Vigilato et al., 2013). SIRVERA is made up of more than 32,000 records, most of which are bovine in origin. Of 29 countries in the Americas in the system, the largest number of reports originate from Brazil, Mexico and Peru. The data was filtered according to the following criteria: **1)** types of cases: animal cases, **2)** date of notification: January 1970 to December 2020, **3)** variant: genetic variant three of the rabies virus (specific to *D. rotundus*), **4)** species: bovine (species of domestic animal most affected), and **5)** aggressor species: sanguivorous bat (refers to the common vampire bat or *D. rotundus*). Rabies outbreaks, difference, and percentage change between the first decade (1970s) and the last decade (2010s) of the data year range were estimated. Additionally, spatial analyzes were performed using classification schemes to visualize area patterns (Moore & Carpenter, 1999; Morgan, Wallace, Vokaty, Seetahal, & Nakazawa, 2020; Nihal, Dangolla, Hettiarachchi, Abeynayake, & Stephen, 2019). The scale or intensity of colors was proportional to the number of cases in choropleth maps (Alegria-Moran, Miranda, Barnard, Parra, & Lapierre, 2017; Blanton, Palmer, & Rupprecht, 2010; Moore & Carpenter, 1999). Choropleth maps were made at the country scale between Mexico and Argentina and Chile (historical distribution area of *D. rotundus*) on the increase in rabies outbreaks between 1970 and 2020. The *dplyr*, *sp*, *rgdal* and *ggplot2* packages of the software R (version 4.1.0) and RStudio (version 2022.02.3) were used.

#### 2.0 Supplementary Methods 2

To assess the impact of retrospective climate change on the distribution of *Desmodus rotundus* we utilized methods from ecological niche modeling to complete a multidimensional modeling effort of the species range across the last century using R statistical software and the Maxent algorithm (J. Phillips et al., 2019) (Supplementary Figure 1). Maxent was utilized because, as a presence-background modeling algorithm, it does not require absence data, and therefore more logically abides by the available occurrence data for this species. Maxent functions by contrasting the environmental conditions associated with presences points (occurrence points) with randomly selected background points from the available environmental space where the species could potentially occur (i.e., study area extent) (J. Phillips et al., 2019; S. Phillips, 2010; S. J. Phillips et al., 2006).

##### 2.1 Climate Data

For background environmental variables, we used six representative climate variables from the Climatic Research Unit gridded Time Series (CRU TS) version 4.04 database at 0.5° latitude by 0.5° longitude resolution from 1901-2019 (Harris et al., 2020). To reduce dimensionality in the final modeling effort and to summarize annual climatic variability we collated the available monthly bioclimatic variables from CRU TS into annual level rasters. Representative climate variable included: average temperature, temperature standard deviation, average diurnal temperature range, average cloud cover, cloud cover standard deviation, and potential evapotranspiration standard deviation. These variables were converted to ASCII format and were used at the annual level for the occurrence data filtering, model calibration, and model projection process.

#### 2.2 Species Occurrence Records and Filtering

Occurrence records of the common vampire bat (*Desmodus rotundus)* were collected from a variety of sources including: 1) publicly available resources and databases, 2) a network of natural history museums across North, Central, and South America, 3) official repositories in ministries of agriculture and health, and 4) from published scientific literature. These data were curated and published in a publicly available data repository(Van de Vuurst et al., 2021). To address possible sample selection bias and spatial autocorrelation we resampled the *D. rotundus* occurrences to one per pixel of the study extent. We then used the remaining occurrence data and the environmental background data to filter the occurrences by environment to identify outliers. The values of each climatic variable were first extracted from the location and year of each occurrence to create a cloud of data points representing the species distribution in environmental space (Supplementary Figure 2) (Peterson et al., 2011). For occurrence records which had age or life stage metadata, we extracted the variable values from multiple years based upon the age of the individual. For juvenile individuals only the singular year of occurrence was extracted from the corresponding annual raster. For individuals that were classified as adults, we extracted the year the occurrence was recorded and the four years prior (five years total). Age delineations were based on previous *D. rotundus* capture data (Lord et al., 1976), which indicates that most captured adult *D. rotundus* individuals are less than six years of age.

We then developed a principal component analysis of the climatic variables to obtain principal component axes which summarized the variance of the data. Principal components 1-3 (summarizing 82.9% of the data variance) were then used as axes (i.e., X, Y, and Z axes) to plot the species occurrence environmental data. We used a minimum volume ellipsoid to identify outliers from the cloud of extracted values, which were removed from the analysis

(Supplementary Figure 2). The remaining filtered occurrences were randomly split into 50% training 50% testing subsets from the thinned dataset.

#### 2.3 Model Calibration and Evaluation

For this modeling effort we used a presence-background ecological niche modeling approach based on maximum entropy (Maxent v3.4.4) (J. Phillips et al., 2019; Warren & Seifert, 2011) within the ***kuenm*** package in R software (Cobos et al., 2019). **kuenm** is a cutting edge R package designed to make the process of model calibration and final model creation more reproducible and robust (Cobos et al., 2019). Using this package, we created suites of candidate models with various parameterizations and regularizations. Model overfitting is minimized in Maxent by the use of functions derived from the environmental variables (i.e. feature classes) and regularization parameters which impose penalties on the model for over-complexity (Morales et al., 2017; S. J. Phillips & Dudík, 2008). We tested a suite of regularization parameters (0.1-1.0 by increasing values of 0.1, 2-6 by values of 1, 8, and 10) and all twenty-nine possible combinations of five feature classes (linear=l, quadratic=q, product=p, threshold=t, and hinge=h) to ensure all possible models were considered. After redundant model combinations were removed, this resulted in 1,054 candidate models for each replication. The candidate model formation, evaluation, and best model selection process was repeated 100 times (for a total of 105,400 models) to increase confidence in the resulting best model features validity via statistical replication. The best model from each replicate was reported and its regularization multipliers and feature classes were recorded. The best candidate model regularization multiplier and feature classes that appeared most often from the 100 replicates were then selected for the final model projection process.

Candidate models were evaluated using the *kuenm_ceval* function. Candidate model performance evaluation was based on significance (partial ROC, with 100 iterations and 80 percent of data for K-fold validation), omission rates (*E*=5%), and model complexity and fit to the calibration data (i.e., Akaike information criterion) (Hobbs & Hilborn, 2006). Both partial ROC and omission rate were used as preliminary measures to identify models which were significantly better than random. AICc was used as the delineating value of best model selection, as AICc is a more meaningful measure of model performance (Lobo et al., 2008; Peterson et al., 2008; Warren & Seifert, 2011). The final model was then used to project *D. rotundus* range across the entire temporal extent of the study (1901-2019).

**2.4 Range Shift Analysis**

The minimum training presence value from the final model was used as a threshold to reclassify the projected range maps into annual binary maps. Based on the assumption that the least suitable environment at which the species is known to occur is the minimum suitability value for the species, this threshold can be used to delineate areas of possible species range from areas where it is unlikely for the species to occur. The resulting binary maps were used to assess how the projected suitable range for *D. rotundus* has changed across time. We used the *cellStats* function of the ***raster*** package (Etten et al., 2021) in R to quantify the total range area from each binary map. We found no significant changes in the total range areas of *D. rotundus* during the 1901-2019 period. We then isolated the 100 highest (most northern) and lowest (most southern) latitudes predicted by the projected range models, which allowed us to assess whether or not the projected suitable range for *D. rotundus* was moving northward or southward across time based upon our model (Figure 3 main article). We also identified the 100 highest projected suitable elevations from each annual binary map and averaged these values across time to determine the extent to which *D. rotundus*' suitable range varied in elevation. Elevation data was collected from the WorldClim bioclimatic variable database at five arc minute resolution (approximately 10km) in raw agreement with the climate data (Fick & Hijmans, 2017). We found no significant trend of elevational change for the species' range. We used linear models to assess the relationships between time as measured by year and the three resulting suitability characteristics (i.e., area, elevation, and latitude of occurrence and predicted range).

To identify if there were trends in species occurrence beyond the model, we isolated the most northern and most southern extents of the species' observed occurrence (i.e., from the occurrence points). We then used a linear mode in the ***stats*** package to identify the relationship between the species occurrence at the extent of its range and time as measured by year. This analysis was limited to the last 50 years due to limited occurrence data availability in the earlier decades of the 1900's. To identify northern range shift rate across time for *D. rotundus* we identified the top 20 most northern occurrences of the species for each year using the ***dplyr*** package in R. This allowed us to ascertain the average most northern extent of the species range for each year based on recorded occurrence. We then calculated the average change in latitude by decimal degrees per year. We then converted this metric to kilometers using the assumption that one decimal degree is approximately 111 km.

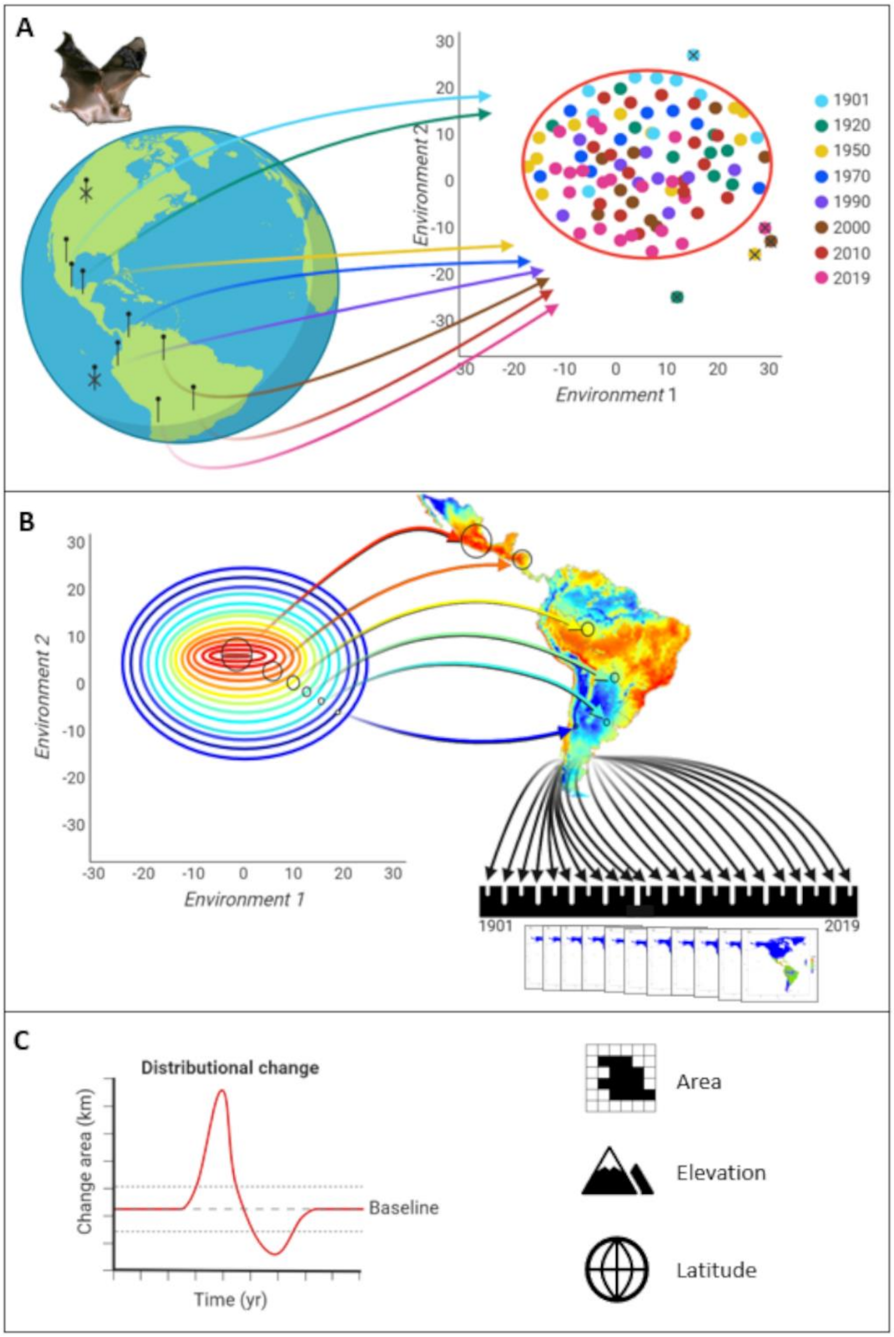


**Supplementary Figure 1: Modeling methods framework. (A)** Occurrence records of *D. rotundus* were collected from a variety of sources[3]. We then filtered the records both by geography (physical location), and by environment. (**B)** For background climate data we used six optimal variables from the Climatic Research Unit gridded Time Series (CRU TS) version 4.04 database[4]. We then utilized the modeling algorithm Maxent to create maps of vampire bat ranges for each year from 1901-2019, (**C)** which we then used to assess how vampire bat ranges had changed in area, elevation, and latitude.

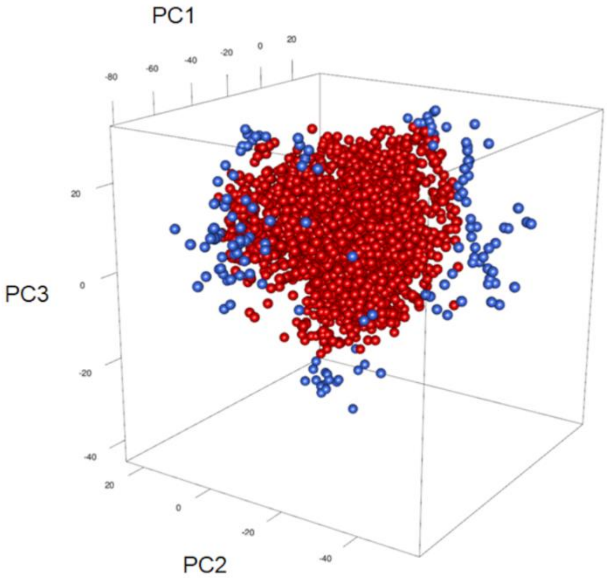


**Supplementary Figure 2: Occurrence filtration in environmental space.** The first three principal components (summarizing 82.9% of the data variance) of the climate variables (average temperature, temperature standard deviation, average diurnal temperature range, average cloud cover, cloud cover standard deviation, and potential evapotranspiration standard deviation) from the CRU TS[4] database at the annual level were used as axes (i.e., X, Y, and Z axes) to plot *Desmodus rotundus* occurrence reports in the resultant three-dimensional environmental space. We used a minimum volume ellipsoid to identify occurrence points that fell outside of this ellipsoid (blue) as outliers, which were removed. Occurrences which were not environmental outliers (red) were retained and used for model calibration.
